# Accelerated Discovery of Aptamer Beacons via Massively Parallel Screening

**DOI:** 10.1101/2025.07.31.667975

**Authors:** Yasser Gidi, Linus A. Hein, Hajime Fujita, Michael Eisenstein, Hyongsok Tom Soh

## Abstract

Aptamer beacons are unimolecular probes that undergo a reversible conformational change upon target binding, making them a promising tool for the real-time detection and monitoring of molecular analytes. However, the development of such sensors has been impeded by the lack of generalizable tools for the efficient discovery and optimization of aptamer beacons for diverse molhhtmlecular targets. Here, we present a scalable approach for converting existing strand-displacement aptamer switches into aptamer beacons by introducing targeted mismatches within their non-target-binding stem domain, destabilizing the aptamer structure to an extent that it can only refold upon binding its target. In order to perform this screening in a high-throughput fashion, we have developed the Massively-parallel Aptamer Performance Analyzer (MAPA), an automated, fluorescence-based screening platform based on a reconfigured sequencing instrument that can functionally evaluate millions of aptamer variants in parallel. Using MAPA, we identified multiple aptamer beacons for glucose, serotonin, and dopamine, and demonstrated that these beacons retain their sensing performance when translated from the on-chip surface-based MAPA format to a solution-based assay. We performed all three aptamer beacon experiments on a single MAPA chip, and even greater multiplexing should be possible, greatly accelerating the discovery of aptamer-based sensors for real-time molecular detection.

## Introduction

Aptamers are useful receptors for real-time biomarker sensing because they can undergo reversible conformational switching in response to specific target-binding events. Researchers have employed a variety of sensing strategies to transduce target-induced conformational changes in aptamers into measurable signals. These include electrochemical aptamer-based sensors, which detect changes in electrical current resulting from structural shifts in surface-bound, redox-tagged aptamers.^1^ Field-effect transistor (FET)-based sensors leverage the movement of negatively-charged phosphodiester backbones toward the FET channel surface to induce electrical signal changes.^2^ A number of groups have also developed optical aptamer-based sensors, where target binding alters the distance between a fluorophore and a quencher to modulate fluorescence.^3^ We and others have used such sensors to achieve sensitive, specific, continuous biosensing in both *in vitro* and *in vivo* settings.^4, 5^

Many aptamer-based sensors employ a strand-displacement aptamer switch design. Here, the aptamer is hybridized to a short, complementary displacement strand labeled with a quencher, while the aptamer itself is tagged with a fluorophore. In the absence of the target, the aptamer and displacement strand form a duplex, bringing the signaling moieties into close proximity and resulting in quenched fluorescence. Upon target binding, the aptamer undergoes a conformational change that leads to the release of the displacement strand, producing a fluorescence increase. Strand-displacement aptamers are commonly generated through capture-SELEX, a reliable and well-established selection method,^6, 7^ and can deliver excellent target specificity. However, because the displacement strand is released during target binding, this design is incompatible with continuous monitoring in flow systems in which the released displacement strand is washed away, rendering the system irreversible.

For such continuous monitoring applications, aptamer beacons offer a promising alternative. These are hairpin-shaped aptamers that are typically labeled with a fluorophore at one end and a quencher at the other, generating a change in fluorescent readout after undergoing binding-induced refolding.^8, 9^ In contrast to bimolecular strand-displacement sensors, aptamer beacons are unimolecular constructs that can reversibly fold and unfold, making them particularly well-suited for continuous monitoring applications. Aptamer beacons have also proven highly effective in electrochemical sensing platforms, in which the aptamer is immobilized onto a surface at one terminus and then functionalized with a redox label at the other. However, there are currently no aptamer selection strategies that are specifically designed for the selection of aptamer beacons,^9^ and the adaptation of existing aptamers into beacons remains a major bottleneck for sensing new targets.^1^

Strategies to engineer aptamer beacons often involve truncation of the stem of native hairpin-or Y-shaped aptamers, with the intention of destabilizing the structure in the absence of the target while enabling hybridization upon target binding.^10-13^ However, stem truncation may also compromise functionality if essential bases are unwittingly removed. Additionally, reducing stem length can diminish the signal change by limiting the magnitude of the end-to-end distance change that occurs between the labeled termini of the beacon. For fluorescent signaling, achieving a 50% change in signal typically requires a distance change exceeding the Förster radius of the fluorophore-quencher pair—usually around 6 nm (equivalent to ∽18 base-pairs of double-stranded DNA).^14^ For electrochemical measurements, the electron transfer rate scales inversely with oligonucleotide length.^15^ In both cases, signal change is expected to decrease by truncating the length of the DNA stem. Furthermore, for a given aptamer, although many variants are capable of signal transduction, the magnitude of signal gain upon target binding can vary by more than two orders of magnitude. For example, the signal gain from ATP sensors in one study varied from -10– 200%.^16^ Therefore, the switching architecture must also be carefully optimized based on the unique structural and functional properties of each aptamer sequence—features that are often poorly understood.

In this work, we present a molecular framework for converting strand-displacement aptamers into aptamer beacons. Our strategy entails the introduction of targeted mismatches into the hybridization region of the strand-displacement aptamer, and then screening for those sequences which remain unfolded in the absence of target but are able to hybridize into their folded form in the presence of target. We have developed the Massively-parallel Aptamer Performance Analyzer (MAPA), a fully automated high-throughput instrumental platform that allows us to perform fluorescence-based screening of large numbers (∽10^6^–10^7^) of aptamer variants, enabling the large-scale exploration of sequence-function relationships and rapid prototyping of aptamer beacons. Using this approach, we successfully identified the top-performing aptamers beacons variants for glucose, serotonin, and dopamine. We then synthesized the highest-ranking candidates, and confirmed that they all exhibited target-responsive fluorescence signals. Notably, their experimental performance correlated closely with the measurements obtained using MAPA, validating the predictive power of our platform and the functionality of the resulting molecular switches.

## Results and Discussion

### Overview of MAPA

To obtain structure-switching capability without the need for a detaching displacement strand, we hypothesized that we could convert existing strand-displacement aptamers (**Figure 1a, left**) into aptamer beacons by introducing mismatches into the stem region that would normally bind to the displacement strand. This would weaken intramolecular hybridization, favoring an unhybridized conformation in the absence of the target (**Figure 1a, right**). In the presence of the target, binding would create energetically-favorable conditions for the mismatched strands to hybridize, thus restoring the aptamer’s reversible switching capability. This requires the mutations to be carefully tuned to favor a moderately destabilized and thus unhybridized conformation in the absence of target while still enabling refolding upon target binding. In most cases, little is known about the specific target interactions or binding mechanisms of the aptamers, making it difficult to identify which specific bases can be mutated without compromising functionality. Our approach therefore involves introducing mutations into regions that were originally involved in hybridization with the displacement strand during the capture-SELEX process (*i*.*e*., the 5’ side of the stem); such alterations will have no impact on stem length and should not directly affect aptamer-target interactions, thus preserving binding function.

**Figure 1.**
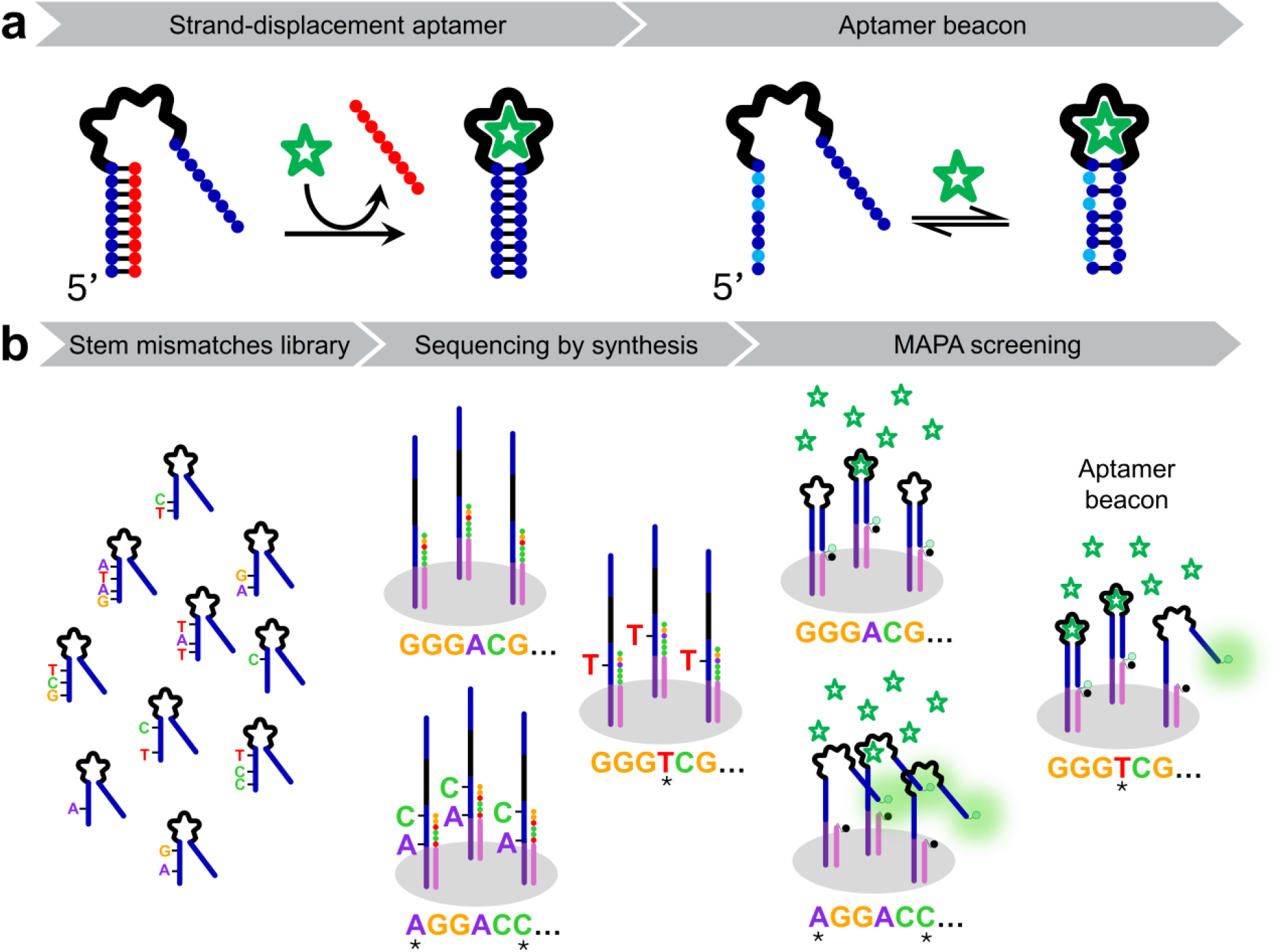
Molecular design and method for converting strand-displacement aptamers into aptamer beacons. **a**. (left) Representation of the strand-displacement aptamer with the displacement strand shown in red. (right) Strand-displacement aptamers are mutated close to the 5’ end of the stem region allowing to remove the displacement strand from the equilibrium to render aptamer beacons. **b**. (left) Oligonucleotide pools were generated with targeted mutations in the 5′-end portion of the stem region of the original strand displacement aptamer. (middle) Libraries were sequenced using High-Throughput sequencing-by-synthesis on a MiSeq platform. (right) The sequenced flow cell was transferred to the MAPA system for high-throughput functional screening to identify aptamer beacons.

In order to maximize the number of possible mutants that we could screen in a single experiment, and thereby increase our odds of a successful screen, we devised a high-throughput platform to efficiently evaluate and select functional variants. Inspired by the work of the Greenleaf lab,^17^ we developed MAPA, in which a Illumina Genome Analyzer IIx (GA_IIx_) sequencer is repurposed for fluorescence-based aptamer screening (**Figure 1b, right;** for a detailed description, see **Supplementary Information, Section i.b and i.c**). The MAPA platform exploits the capabilities of a high-throughput DNA sequencing instrument to achieve parallel functional analysis of millions of DNA clusters. Our choice to use GA_IIx_ as the foundation for MAPA was driven by its internal laser-illuminated, prism-based total internal reflection fluorescence (TIRF) imaging system, which is highly effective at minimizing background signal. This TIRF architecture also enables future adaptation to other assays involving fluorescently-labeled targets, expanding the platform’s versatility. This high sensitivity of TIRF is highlighted by its widespread use in demanding applications such as single-molecule fluorescence spectroscopy.^18^ This is in contrast to our previously developed N2A2 platform, which employs a modified MiSeq instrument. That system was optimized for color-based base-calling rather than precise signal quantification, making it challenging to reliably extract accurate binding affinity and kinetic parameters from an N2A2 experiment.

For this set of MAPA screening experiments, we began by generating large, mutagenized libraries targeting the 5’ end of the double-stranded stem region of an existing strand-displacement aptamer (**Figure 1b, left**). In principle, any aptamers obtained via capture-SELEX can be subject to this screening. These libraries were then sequenced via sequencing-by-synthesis using a MiSeq instrument. (**Figure 1b, middle**), during which each aptamer candidate became immobilized as a DNA cluster on the flow cell surface, with each cluster comprising many copies of a single sequence. After modifying the aptamers with fluorescent moieties, the flow cell was exposed to the target molecule and imaged to quantify aptamer fluorescence response in parallel across millions of clusters (**Figure 1b, right**). The GA_IIx_ sequencing instrument itself was reconfigured to accommodate a MiSeq flow cell mounted over a prism and illuminated with a 532-nm laser (**Figure 2**). Laser light was coupled into a multimode fiber optic and routed through an optical switch for precise control of illumination. The output was then passed through a multimode scrambler to improve illumination homogeneity before being directed in free space through the prism and onto the flow cell surface. Fluorescence emission was collected through an air-immersion objective, passed through a tube lens, and projected onto a camera chip. An emission filter mounted in a motorized filter wheel was positioned between the tube lens and the camera to reject elastically-scattered laser light and isolate the desired spectral fluorescence band. The flow cell-prism assembly was mounted on a motorized XY stage, while the objective was mounted on a motorized Z stage. Temperature was maintained by a thermoelectric cooler device placed beneath the prism, as previously reported.^17^ Temperature was monitored via thermocouples and controlled through a closed-loop feedback system. Reagents were delivered using a syringe pump that draws fluids through the flow cell, with multiple reagents accessed via a multiport valve. Since the original camera from the GA_IIx_ instrument lacked sufficient coverage for the wide imaging area required to capture the entire MiSeq flow cell, we integrated a large-chip Teledyne Kinetix sCMOS camera, as this camera also supports high-speed kinetic measurements and offers improved quantum efficiency. Finally, to achieve fully-automated operation, we implemented an unsupervised focus-finding system and developed custom software for complete computer control and workflow automation.

**Figure 2.**
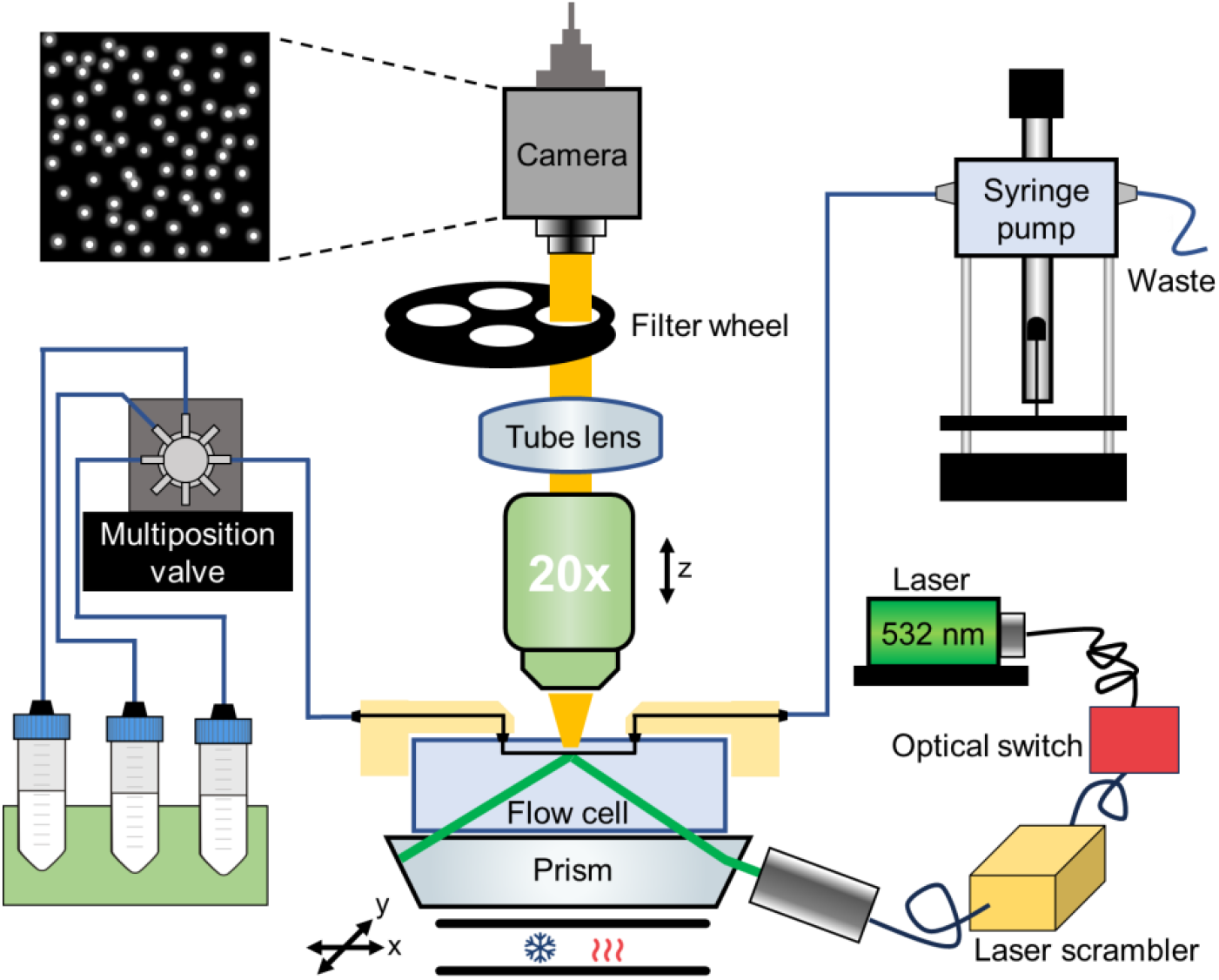
Massively-parallel aptamer performance analyzer (MAPA) hardware for fluorescence-based assays. A GA_IIx_ sequencing instrument was modified to accommodate a MiSeq flow cell illuminated by a 532 nm laser. Emitted fluorescence was collected through an air-immersion objective, passed through a motorized emission filter, and imaged by a large-area sCMOS camera. The flow cell was mounted on a motorized XY stage, with temperature control. Reagents were delivered through a syringe pump and multiport valve. Custom software enables fully automated assay workflows.

### Screening methodology

We first designed a mutant library to convert existing strand-displacement aptamers into aptamer beacons and commissioned its synthesis from TWIST Biosciences. The library was based on previously-reported strand-displacement aptamers for glucose (GLU-SD, K_d_: 150 mM), serotonin (SER-SD, K_d_: 0.4 μM), and dopamine (DOPA-SD, K_d_: 10 μM) (**Figure 3a**).^2^ To guide library mutagenesis, we predicted the secondary structures of each original aptamer using MFold.^19^ **Figure 3a** displays the most stable predicted structure for each aptamer, though we considered the three next most thermodynamically-favorable structures in our design as well (**Supplementary Figures s3-5**).

**Figure 3.**
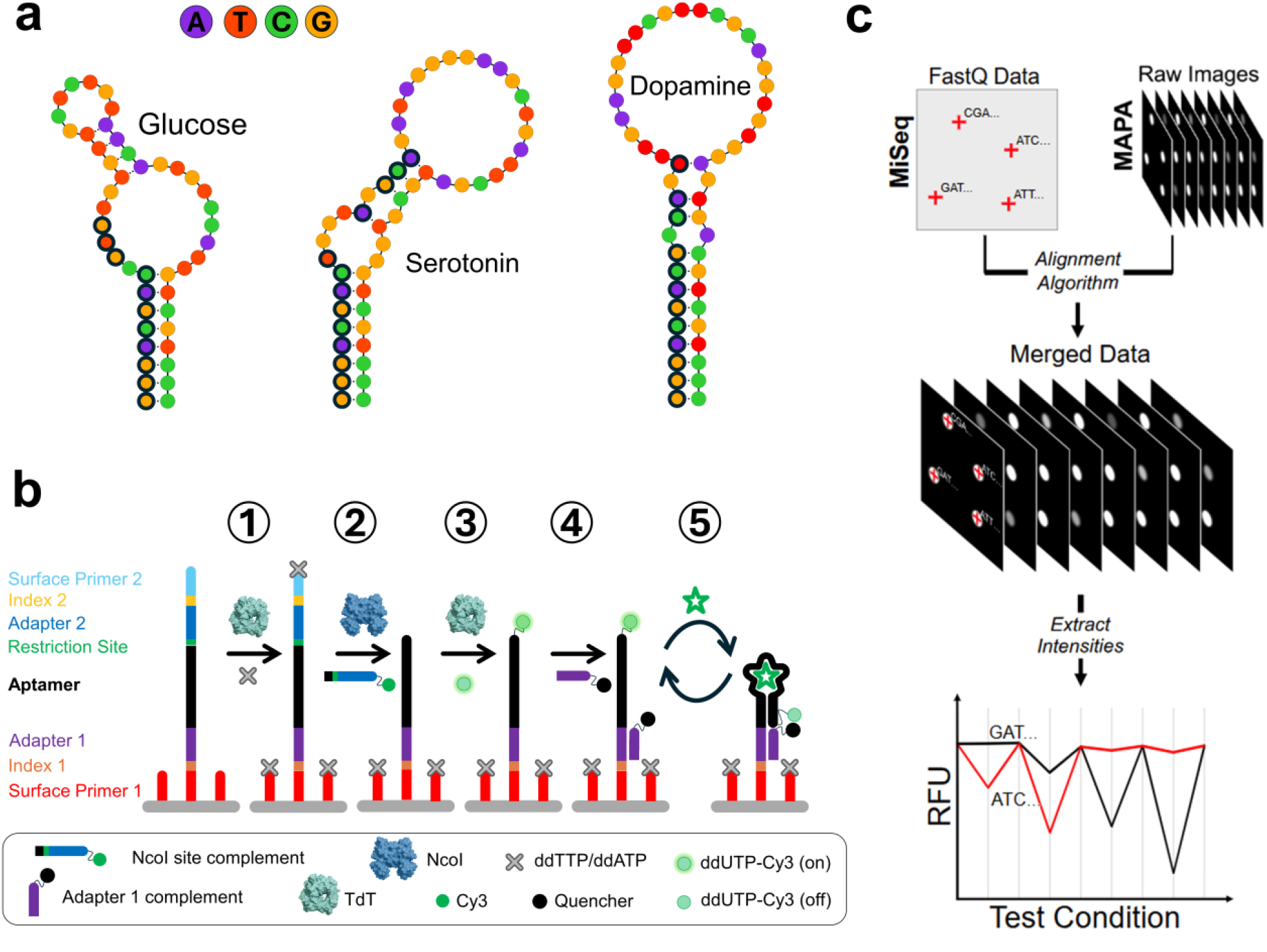
MAPA screening methodology. **a**. Predicted secondary structure of the three strand-displacement aptamers studied in this work, generated using Mfold. Bases selected for mutagenesis are outlined in thick black. **b**. Post-sequencing cluster preparation. Clusters were processed to enable target-induced signal measurements through the following steps. (1) Flow cell lawns and DNA clusters were blocked using terminal deoxynucleotidyl transferase (TdT) and ddNTPs. (2) Sequencing-specific regions were cleaved away using the restriction enzyme *NcoI*. (3) Cleaved clusters were fluorescently labeled using TdT and ddUTP-Cy3. (4) A quencher-labeled strand (Dabcyl) complementary to sequencing adapter 1 was hybridized. (5) The flow cell was exposed to varying concentrations of target in buffer, interspersed with target-free buffer solutions to assess reversibility and identify target-responsive aptamer beacons. **c**. Data analysis involved coordinate alignment between the MiSeq and MAPA imaging systems, extraction of fluorescence signals for each cluster, and data visualization to identify functional aptamer beacons.

We specifically focused on bases near the 5′ end that participate in canonical base-pairing for mutagenesis, selecting 11 positions for GLU-SD, 13 for SER-SD, and 14 for DOPA-SD (outlined in thick black in **Figure 3a**). We systematically generated all possible single- and double-mutants at these positions. For higher-order variants containing 3–6 mutations, we randomly selected the identity of the replacement nucleotide at each mutated position. This strategy enabled broad coverage of combinatorial sequence space while maintaining a tractable library size, although it did not exhaustively sample all possible substitutions at each site. Using this approach, we constructed mutant libraries containing 1,948, 4,746, and 7,232 aptamer variants for glucose, serotonin, and dopamine, respectively. These aptamer variant sequences were flanked by Illumina Nextera Transposase adapters sequences, with adapter 1 at the 5’end and adapter 2 at the 3’end (**Figure 3b**). An *NcoI* restriction site (C/CATGG) was inserted between the aptamer and adapter 2. We selected *NcoI* because its recognition sequence begins with a cytosine, enabling us to match the final base of all aptamers tested —ensuring that after enzymatic cleavage, we are left with the exact original aptamer sequence without any extra or missing nucleotides. Finally, these library molecules were flanked by indices and surface primers via limited cycle PCR and then sequenced on the MiSeq.

After sequencing, the flow cell was transferred to the MAPA platform, where the DNA clusters underwent a four-step molecular pre-processing procedure (**Figure 3b; SI section i.h**). First, we blocked the 3’ end of both the clusters and surface primers using a mixture of ddATP, ddTTP, and terminal deoxynucleotidyl transferase (TdT). We next hybridized a complementary strand encoding a sequence complementary to the *NcoI* restriction enzyme site to the cluster strands, and then added that restriction enzyme to cleave at the *Ncol* site, removing the DNA regions that were only required for sequencing (*i*.*e*. surface primer 2, index 2, adapter 2, and the *NcoI* restriction site). Next, the cleaved clusters were fluorescently labeled using TdT and ddUTP-Cy3. Finally, we hybridized the cluster strands to a quencher-labeled strand that is complementary to the sequencing ‘adapter 1’ region. We next exposed the flow cell to varying concentrations of target in buffer, interspersed with target-free buffer solutions to assess reversibility and identify target-responsive aptamer beacons. Upon target binding, such aptamers undergo a conformational change that brings the 3′-end Cy3 label into close proximity to the quencher, resulting in a decreased fluorescence signal. For each experimental condition, our MAPA platform captures 19 fluorescence images of the sequencing flow cell at positions matching images taken by the MiSeq platform (**Figure 3c, top right**). Fluorescently-labeled DNA clusters were detected and aligned with the sequencing output files (FastQ), which provide positional and sequence identity information for each cluster (**Figure 3c, top left**). We then extracted the fluorescence intensity of each cluster and aggregated these intensities across all clusters and conditions. To identify functional variants, we plotted the fluorescence of each sequence versus experimental condition to discover mutants that exhibit a robust change in fluorescence in the presence of target (**Figure 3c, bottom, Supporting Information section i.d**). Notably, our MAPA platform enabled the simultaneous screening of GLU-SD, SER-SD, and DOPA-SD within a single flow cell, because cluster families corresponding to each aptamer library can be computationally separated during data analysis. While we have demonstrated this capability with three aptamers, the platform is readily scalable and not limited to this number, allowing for the parallel screening of many more aptamer candidates in future applications.

### Discovering aptamer beacons for multiple small-molecule targets

We next sought to identify aptamer beacons that respond to their respective targets. For each target, we tested a range of concentrations spanning 0.001–100 mM for glucose, and 0.001–100 μM for serotonin and dopamine. For each unique sequence, we averaged the fluorescence intensities across all clusters sharing that sequence. We found an average of 91, 93, and 90 copies representing each aptamer variant for glucose, serotonin, and dopamine, respectively, providing sufficient redundancy for robust quantification. To quantify the response of each variant, we calculated the normalized fluorescence change (Δ*S*_*N*_) using the formula Δ*S*_*N*_ = (*I*_*T*_ − *I*_*b*_)/*I*_*b*_, where *I*_*b*_ and *I*_*T*_ represent the fluorescence intensities measured in buffer and target solutions, respectively. Measurements at the highest target concentration were performed in triplicate, and the resulting responses were averaged and ranked to identify the top-performing sequences (**Figure 4a**).

**Figure 4.**
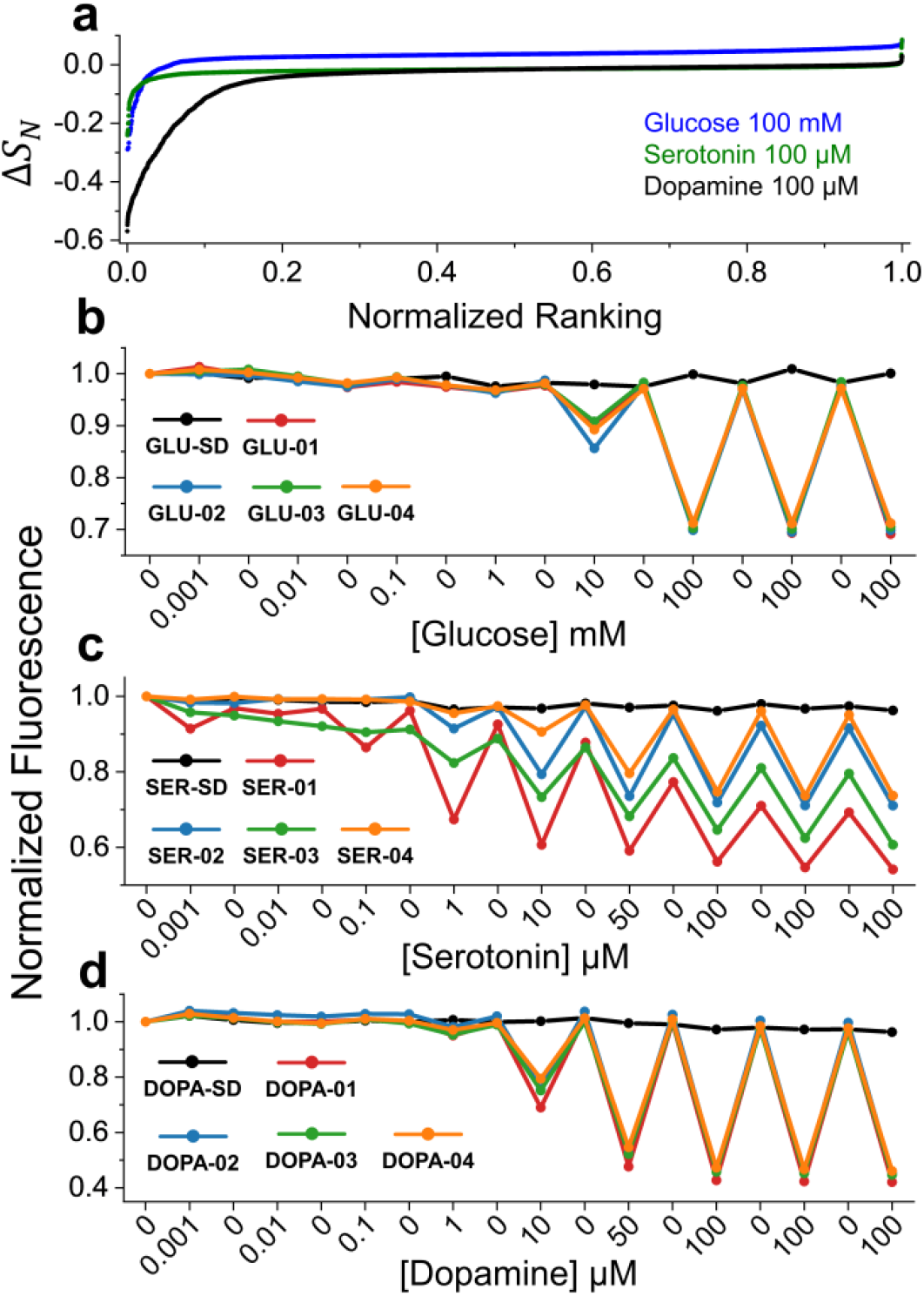
Identification of high-performing aptamer beacons for glucose, serotonin, and dopamine using MAPA. **a**. Normalized signal change as a function of normalized rank for all aptamer variants screened against each target at the highest concentration tested. Only a small fraction of variants display notable target-induced signal changes. **b–d**. Normalized fluorescence responses of the top four MAPA-identified aptamer beacons for glucose (**b**), serotonin (**c**), and dopamine (**d**) compared to their respective parental aptamers (GLU-SD, SER-SD, and DOPA-SD).

Only a small subset of aptamer variants generated a meaningful signal. For glucose, the largest magnitude Δ*S*_*N*_ observed was -0.29, yet only the top 1.4% of variants produced a Δ*S*_*N*_ of ≤ -0.1, underscoring the importance of high-throughput screening to find those rare sequence variants that maximize signal change. For serotonin, the largest Δ*S*_*N*_ was -0.24, with only the top 6.7% of variants meeting the Δ*S*_*N*_ ≤ −0.1 threshold. Dopamine aptamers were most prone to forming aptamer beacons, with a largest Δ*S*_*N*_ = −0.57 and the top 11.1% of variants achieving Δ*S*_*N*_ ≤ −0.1. Interestingly, for glucose, the ΔS_N_ values did not converge toward zero with decreasing rank. For instance, at the 80% percentile, we observed a positive Δ*S*_*N*_ of 0.046, resulting in an upward shift of the curve relative to those for dopamine and serotonin. This behavior may reflect the viscosity-dependent fluorescence properties of Cy3, where fluorescence intensity increases in more viscous environments such as those created by high concentrations (100 mM) of glucose.^14, 20, 21^

Next, we plotted the top four responding aptamer beacons for each target. As expected, the original aptamers GLU-SD, SER-SD, and DOPA-SD were minimally responsive to their respective targets. Only GLU-SD showed a slight increase in fluorescence, consistent with the effect of high glucose concentration proposed above. The consistently low fluorescence intensities observed across all SD aptamers support the notion that these constructs predominantly remain fully-folded, resulting in target-independent quenching of the 3′ Cy3 fluorophore (**Figure S7)**. In contrast, the top four glucose-responsive aptamer beacons identified via MAPA—GLU-01, GLU-02, GLU-03, and GLU-04—exhibited normalized signal changes ranging from -0.29 to -0.26 at the highest tested glucose concentration (100 mM) (**Table 1, Table 2, and Figure 4b**). Similarly, the top serotonin aptamer beacons, SER-01, SER-02, SER-03, and SER-04, showed normalized signal changes between -0.24 and -0.23 at 100 μM serotonin (**Figure 4c**). Interestingly, a mutation at base 17A was the most prevalent among the top-performing variants, appearing in 28 out of the top 30 mutants. Although this position lies far from the 5′ end, it appears to be critical for aptamer beacon functionality—a feature that would have been difficult to predict *a priori*, highlighting the power of our high-throughput mismatch screening approach. We also observed an interesting temporal signal drift with SER-01 and SER-03, which is likely due to slow aptamer binding and unbinding kinetics, where the 10-minute incubation used in the MAPA screen may not have been sufficient to reach maximal signaling in the presence of target or to return to baseline levels during the buffer cycle. The dopamine aptamer beacons demonstrated the strongest signal responses in our MAPA screen. The top variants, DOPA-01–04, exhibited normalized signal changes between -0.57 and - 0.53 at 100 μM dopamine (**Figure 4d**).

**Table 1.**
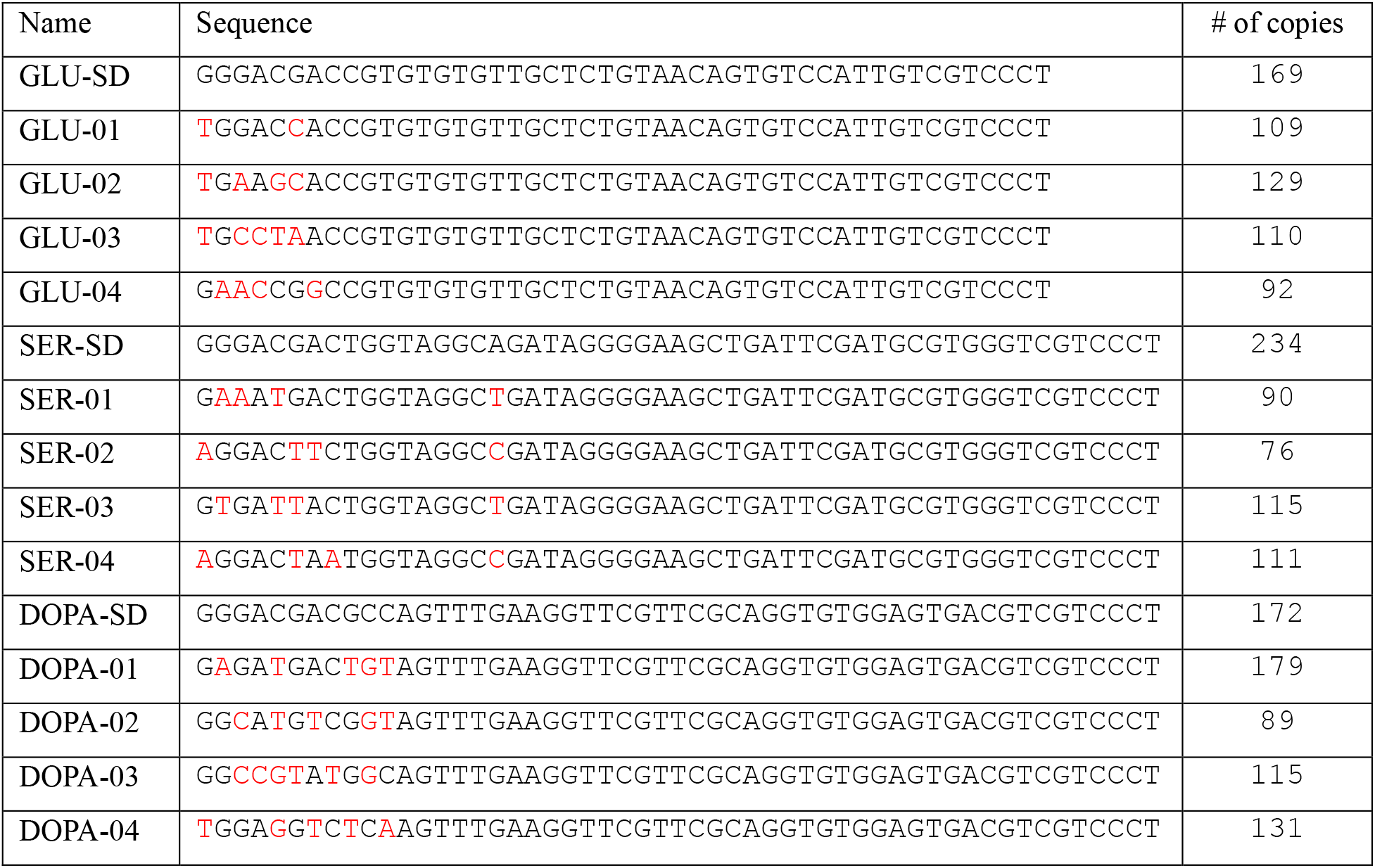
Sequences of original aptamers and MAPA-selected aptamer beacons. Red letters represent mutated bases. All oligos were appended with a 3’-terminal thymine to account for the base added by TdT in MAPA.

**Table 2.** Fluorescence responses of original aptamers and MAPA selected aptamers beacons via MAPA and in bulk solution.

| | Fitting parameters | | $\Delta S_N(@T_{max})$ | | |
| --- | --- | --- | --- | --- | --- |
| Name | $K_D$ | $B_{max}$ | Solution | MAPA | $T_{max}$ |
| GLU-01 | $28 \pm 5$ mM | $0.42 \pm 0.02$ | -0.32 | -0.29 | 100 mM |
| GLU-02 | $42 \pm 4$ mM | $0.37 \pm 0.01$ | -0.27 | -0.29 | |
| GLU-03 | $23 \pm 5$ mM | $0.20 \pm 0.01$ | -0.16 | -0.28 | |
| GLU-04 | $19 \pm 6$ mM | $0.29 \pm 0.02$ | -0.25 | -0.26 | |
| SER-01 | $1.2 \pm 0.1$ μM | $0.55 \pm 0.01$ | -0.56 | -0.24 | 100 μM |
| SER-02 | $64 \pm 12$ μM | $0.42 \pm 0.04$ | -0.26 | -0.24 | |
| SER-03 | $13 \pm 1$ μM | $0.272 \pm 0.003$ | -0.24 | -0.23 | |
| SER-04 | $38 \pm 41$ μM | $0.10 \pm 0.05$ | -0.08 | -0.23 | |
| DOPA-01 | $47 \pm 6$ μM | $0.59 \pm 0.01$ | -0.39 | -0.57 | 100 μM |
| DOPA-02 | $47 \pm 11$ μM | $0.56 \pm 0.02$ | -0.39 | -0.55 | |
| DOPA-03 | $199 \pm 40$ μM | $0.49 \pm 0.02$ | -0.17 | -0.54 | |
| DOPA-04 | $140 \pm 24$ μM | $0.51 \pm 0.02$ | -0.29 | -0.53 | |

### Characterizing the properties of the top MAPA-identified aptamer beacons

To validate the performance of the top-performing aptamer beacons identified by MAPA, we synthesized the top four aptamers for each target. Each aptamer was labeled with a FAM fluorophore at the 5′ end and a Dabcyl quencher at the 3′ end, yielding a signal-off design in which fluorescence decreases upon target binding. For consistency with the MAPA protocol, an additional 3′ terminal thymidine was included in all sequences (**Table 1**). We replaced the 5’ Dabcyl/3’ Cy3 labels used in MAPA with 5’ FAM/3’ Dabcyl labels in order to evaluate whether switching behavior was influenced by the fluorophore-quencher identity or position. The aptamer probes were incubated with their respective targets under conditions matching the MAPA protocol. For glucose and dopamine aptamers, a 10-minute incubation was sufficient, but the serotonin aptamers exhibited slower kinetics and were instead incubated for 12 hours to ensure equilibration was reached.

Each aptamer beacon was evaluated in solution across a range of target concentrations. All constructs show dose-dependent decreases in fluorescence upon target addition, consistent with the target-induced quenching behavior observed in MAPA (**Figure 5a-c**). Data points were fitted using the Hill equation (with a fixed Hill coefficient, n = 1) to extract key parameters such as the dissociation constant (K_d_), and maximum signal change (B_max_) (**Supporting Information section i.i**). Glucose-targeting aptamer beacons showed a relatively narrow K_d_ range of 19–42 mM; this is notably lower than the reported 150 mM K_d_ for GLU-SD, indicating improved responsiveness. Serotonin-targeting aptamer beacons exhibited K_d_ values ranging from 1.2 to 54 μM, with the lower end approaching the affinity of the parental SER-SD aptamer (0.4 μM). Dopamine-targeting aptamers beacons displayed a K_d_ range of 47–199 μM, which is relatively high compared to that of DOPA-SD (10 μM). Together, these results suggest that our strategy to transforming aptamers into beacons does not greatly compromise binding affinity, and in some cases may even improve functional response. Among the glucose-targeting aptamer beacons, GLU-01 showed the largest *B*_*max*_ value (0.42), along with a good affinity (28 mM), a favorable balance between affinity and signal change. Similarly, DOPA-01 and SER-01 exhibited both the highest affinity within their respective panels (47 μM and 1.2 μM) and the largest *B*_*max*_ values (0.59 and 0.55, respectively).

**Figure 5.**
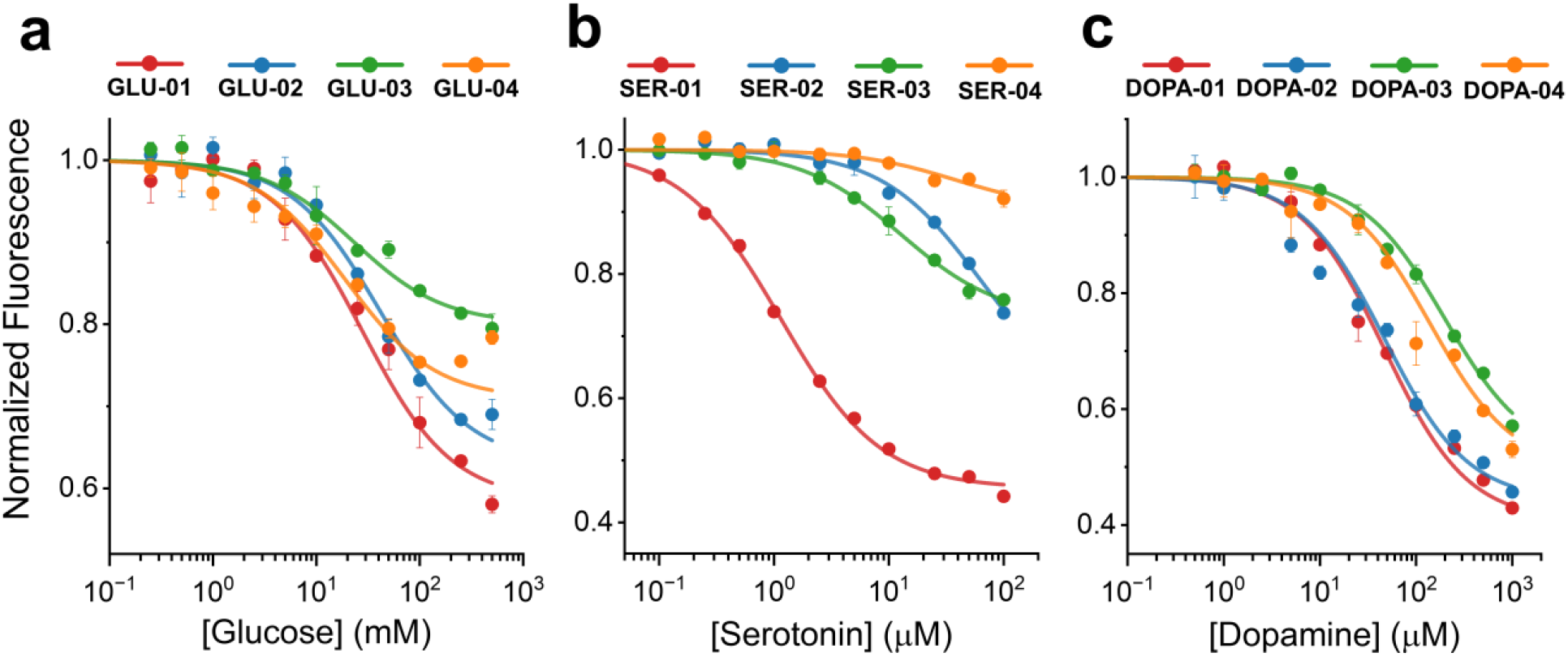
Fluorescence characterization of aptamer beacons. Solution-based fluorescence measurements of the top four MAPA-identified aptamer beacons for **a**. glucose, **b**. serotonin, and **c**. dopamine.

All MAPA-selected sequences exhibited target-responsive signaling in bulk solution assays; however, we observed differences in signal magnitude between platforms. To evaluate cross-platform consistency, we compared the normalized signal change at the maximum target concentration used in MAPA, ΔS_*N*_(@*T*_*max*_) (**see Table 2**). For most glucose beacons, signal responses in solution closely match those measured by MAPA, differing by no more than 0.03 normalized units, except for GLU-03, which showed a 0.12-unit lower response in solution. Results for the serotonin beacons were more varied. SER-02 and SER-03 matched MAPA values closely (within 0.02 units), while SER-01 displayed a significantly higher response in solution (0.32 units above MAPA). This discrepancy likely reflects the slow equilibration kinetics of this aptamer. SER-04, in contrast, showed a 0.15-unit lower response in solution. All dopamine beacons showed decreased Δ*S*_*N*_ values in solution compared to MAPA. These decreases ranged from 0.16 units (DOPA-02) to 0.37 units (DOPA-03), suggesting a family-wide trend.

Several factors could account for these signal deviations, including differences in assay format and fluorophore configuration. In surface-tethered assays (MAPA), aptamer beacons are spatially constrained, whereas in solution they are free to undergo intermolecular interactions, potentially affecting signaling behavior. Additionally, differences in fluorophore identity and labeling position can impact quenching efficiency. Both Cy3/Dabcyl and FAM/Dabcyl pairs support efficient contact-mediated Dexter quenching, but Cy3/Dabcyl exhibits greater Förster resonance energy transfer (FRET) efficiency.^22^ These differences may influence signal magnitude depending on the extent of target-induced distance changes between fluorophore and quencher. This provides a plausible explanation for the family-wide decrease in signaling observed with the dopamine beacons. Notably, the glucose beacons contain relatively short stem regions, whereas the dopamine beacons feature longer stems. Shorter stems are more likely to rely on contact-mediated quenching mechanisms, which are similarly efficient across both fluorophore-quencher pairs. In contrast, longer stems may result in larger distance changes upon target binding, increasing the contribution of FRET-based quenching-where the differences in FRET efficiency between Cy3/Dabcyl and FAM/Dabcyl become more pronounced. Despite these variations, all aptamer beacons retained target-responsive behavior across platforms, supporting the utility of MAPA as a predictive tool for identifying functional aptamer switches.

## Conclusion

In this work, we present a generalizable strategy to generate reversible aptamer beacons for molecular detection by introducing targeted mismatches into the hybridization region of strand-displacement aptamers. This eliminates the need for a separate displacement strand while preserving target-induced structural switching. To enable the high-throughput screening of many such designs, we have developed MAPA: a fully-automated, fluorescence-based screening platform capable of evaluating millions of aptamer variants in parallel. MAPA enables rapid exploration of the vast sequence-function landscape, eliminating the need for low-throughput, trial-and-error optimization approaches. Using this approach, we successfully generated aptamer beacons for glucose, serotonin, and dopamine targets, and demonstrated that the beacons identified in this chip-based assay format retain their performance when employed for solution-phase detection. Our approach is highly scalable and compatible with multiplexing, as it leverages the principles of high-throughput sequencing to enable the simultaneous optimization of multiple aptamer sensors, accelerating progress in the development of aptamer-based sensors for continuous molecular detection.

## Acknowledgments

We thank Nghi Torres for her assistance with custom DNA synthesis.

## Author contributions

Conceptualization: Y.G., H.T.S.

Methodology: Y.G., L.A.H.

Investigation: Y.G., L.A.H., H.F.

Visualization: Y.G., L.A.H.

Funding acquisition: H.T.S.

Project administration: H.T.S.

Writing – original draft: Y.G., H.T.S.

Writing – review & editing: all authors.

## Data and materials availability

All data are available in the main text or the supporting information.

