## Supporting Information for "Accelerated Discovery of Aptamer Beacons via Massively Parallel Screening"

**<u>Table of Contents</u>:**


- **i. Materials and Methods**
  - ***i.a Materials***
    - *Reagents*
    - *Oligonucleotides*
    - *Proteins*
  - ***i.b MAPA Hardware***
    - *Imaging*
    - *Focus-finder*
    - *Fluidics*
    - *Temperature Control*
    - *Motorized Stages*
      - **Figure S1. MAPA hardware configuration**
      - **Figure S2. Focus-finder optical configuration**
      - **Figure S3. Fluidics configuration**
  - ***i.c MAPA Software***
    - *MAPA Software*
    - *Autofocusing*
    - *Event Loop*
    - *Illumination Control*
  - ***i.d MAPA Data Analysis***
    - *Peak Detection*
    - *Coordinate Alignment*
    - *Camera Distortion Correction*
    - *Tile Coordinate Estimation*
    - *Movie Coordinate Alignment*
    - *Intensity Extraction*
    - *Data Aggregation*
  - ***i.e* Library Preparation and High-throughput Sequencing.**
      - **Figure S4. MFold analysis of strand-displacement glucose aptamer**

- **Figure S5. MFold analysis of strand-displacement serotonin aptamer**
- **Figure S6. MFold analysis of strand-displacement dopamine aptamer**

***i.f*** ***Oligo synthesis and purification***

- *Solid-phase Synthesis of Oligonucleotides*
- *HPLC Purification of oligonucleotides*
- *Desalting of Oligonucleotides*

***i.g*** **Oligo Sequences: Designed Oligos for MAPA Screening and Auxiliary Strands**

- **Table S1. Library design**
- **Table S2. Flow cell complementary strands**

***i.h*** **MAPA Molecular Protocols – Cluster Preprocessing**

- *Flow cell Pre-cleaning*
- *Hybridization for Alignment*
- *Lawns and Cluster Blocking*
- *Cluster Cutting*
- *Cluster Labeling*

***i.i*** **Measurement of Effective Binding Affinity via Fluorescence**

***i.j*** **Raw fluorescence responses of the top four MAPA-identified aptamer beacons for glucose, serotonin, and dopamine**

- **Figure S7. Identification of high-performing aptamer beacons for glucose, serotonin, and dopamine using MAPA**

**ii.** **References**

### i. Materials and Methods

#### *i.a Materials*

**Reagents.** 1 M Tris-HCl buffer (pH 7.5) (Cat# BP1757), GelStar (Cat# BMA50535), phosphate-buffered saline (Cat# BP399), ammonium hydroxide (28-30% (w/w) solution in water) (Cat# A669), 1 N NaOH (Cat# SS266-1), 1 M $MgCl_2$ (Cat# AM9530G), Tris(2-carboxyethyl) phosphine (TCEP) (Cat# PG82080), and HPLC-grade acetonitrile (Cat# A998-4) were purchased from Thermo Fisher Scientific. 5 M NaCl (Cat# S5150), anhydrous dimethyl sulfoxide (DMSO) (Cat# 276855), Tween 20 (Cat# P9416), 3 M NaOAc (Cat# 567422), serotonin hydrochloride (Cat# H9523), dopamine hydrochloride (Cat# H8502), and D-(+)-glucose (Cat# G8270) were purchased from Sigma-Aldrich. 1M KCl (Cat# P035) and 1M $CaCl_2$ (Cat# C0477) were purchased from Teknova. HyPure molecular biology-grade water (Cat# SH31191) was acquired from Cytiva. Immersion Liquid (Cat# 19570, Code# 50350, n = 1.4730) was purchased from Cargille. Cy3-labeled ddUTP (5-propargylamino-ddUTP-Cy3; Cat# NU-1619-CY3), unlabeled ddTTP (2',3'-dideoxythymidine-5'-triphosphate; Cat# NU-1018L), and unlabeled ddATP (2',3'-dideoxyadenosine-5'-triphosphate; Cat# NU-1015L) were ordered from Jena Bioscience. 6-Fluorescein phosphoramidite (Cat# 10-1964), 3'-Dabcyl CPG (Cat# 20-5912) and standard phosphoramidites and oligonucleotide synthesis reagents were purchased from Glen Research. All reagents for DNA sequencing were obtained from Illumina. Dabcyl CED phosphoramidite (Cat# CLP-1522) was purchased from Chemgenes.

**Oligonucleotides.** Oligonucleotides shown in **Table S2** (except for FAc-Dabcyl) were purchased from Integrated DNA Technologies (IDT). Labeled and unlabeled oligos were ordered with HPLC and PAGE purification, respectively. Oligonucleotides shown in **Table S3-S5** as well as FAc-Dabcyl (**Table S2**) were synthesized and purified as specified in **section i,e**. All oligonucleotides were resuspended in nuclease-free water and stored at −20 °C.

**Proteins.** NcoI-HF (Cat# R3193L) and terminal deoxynucleotidyl transferase (TdT) (Cat# M0315L) were purchased from New England Biolabs. Terminal transferase, recombinant (Cat# 03333574001) was purchased from ROCHE. GoTaq® G2 Hot Start Master Mix (Cat# M7433) was purchased from Promega.

### *i.b* MAPA Hardware

The MAPA hardware system was developed by repurposing components from an Illumina Genome Analyzer IIx ($GA_{IIx}$) DNA sequencing platform combined with other commercially available components.

**Imaging.** Fluorescence imaging was carried out in an upright prism-based total internal reflection fluorescence (TIRF) microscope configuration using an air-immersion objective (CFI Plan Apochromat Lambda D 20× objective lens, N.A. 0.80, W.D. 0.80 mm, F.O.V. 25 mm, Nikon). To induce total internal reflection at the interface between the flow cell and the internal aqueous medium, a laser beam is directed through a prism onto the flow cell. The imaging platform was equipped with a 532-nm laser (Gem 532, Laser Quantum, $GA_{IIx}$) for excitation of Cy3 and ATTO-550. Although not used in this study, a 660-nm laser (85-RCA-400, Melles Griot, $GA_{IIx}$) was also available for excitation of red-absorbing dyes. Both laser beams were coupled into a multimode fiber optic (GAIIx) and routed through a computer-controlled optical switch (COR ALIGN ILCO12, Luminos Photonics, $GA_{IIx}$). To improve illumination homogeneity, we used a de-speckler (FC to FC - 50 µm core fiber, DSB-0309, Fiberguide) and a dynamic multimode scrambler (MMS-101B-6X-ILM, General Photonics, $GA_{IIx}$), which were connected serially. Emitted fluorescence was spectrally filtered with two emission filters (ET542lp, ET585/65m, Chroma Technology) mounted on a motorized filter wheel (ASI FW-1000-BR, $GA_{IIx}$). All images were recorded onto a 3200 × 3200-pixel region of a back-illuminated scientific CMOS camera (Kinetix, 01-KINETIX-M-C, 1.44 MP, Teledyne). An Arduino UNO was used to synchronize the activation of the laser switch with the camera, compensating for the 5-ms delay inherent in the laser switch.

**Focus-finder.** To achieve unsupervised focusing for automated operation, a fiber-coupled 635-nm laser diode (LPS-635-FC, 2.5 mW, Thorlabs) was collimated and launched into the objective to hit the surface at an angle (**Figure S1**). The reflected beam position in the images, analyzed by a custom algorithm, provides precise information about objective-to-surface distance, requiring only a single initial calibration. This autofocus laser is regulated by a computer-controlled driver (K-Cube KDL101, Thorlabs).

**Fluidics.** Solutions were drawn through the flow cell using a syringe pump (VersaPump 6 8-Channel, 48k, 24741, Kloehn, $GA_{IIx}$). To automatically select the source of liquid reagents, a 26-port multiposition valve (EMHMA-CE, VICI, GAIIx) was employed. Additionally, two automated four-port, two-position selector valves (uProcess AV202-T116, LabSmith) were incorporated into the configuration as shown in **Figure S2**. This arrangement offers two key advantages: (1) the ability to bypass the flow cell (for example, during priming of tubing), and (2) the ability to introduce a new reagent line near the flow cell (Reagent 0). The latter function is particularly critical in two scenarios. The first is when conducting experiments that require rapid changes in reagent concentration. We observed that the dead volume of the VICI multiposition valve produces a concentration gradient in fluids entering the flow cell, resulting in a delay before the fluid reaches its maximum intended concentration. The second scenario is when using costly reagents, as it minimizes the dead volume.

**Temperature Control.** A bipolar (heating and cooling) proportional-integral-derivative temperature controller (TC-36-25-RS232-UL, TE Technology) and thermoelectric module located under the prism (VT-127-1.0-1.3-71P, TE Technology) were used to control the flow cell temperature. Two thermistors (MP-2444, TE Technology)—one located under the prism and the other on the side of the prism—were used to track the temperature of the flow cell surroundings. A recirculating chiller kept at 15 °C was used as a heat sink (UC160, 10-160-G3-P1, Solid State Cooling Systems). The chiller recirculates high-performance liquid coolant (LIQ-702YL-B, Koolance)

**Motorized Stages.** An x-y motorized stage (MS 2000 XY stage, Applied Scientific Instrumentation (ASI), $GA_{IIx}$) was used to position the flow cell under the objective mounted in a z motorized stage (LS50A, ASI, $GA_{IIx}$). Motorized x-y and z stages as well as a filter wheel were controlled by a RS232 controller (LX-4000, LX-XYB-Z5A-FW, ASI, $GA_{IIx}$).

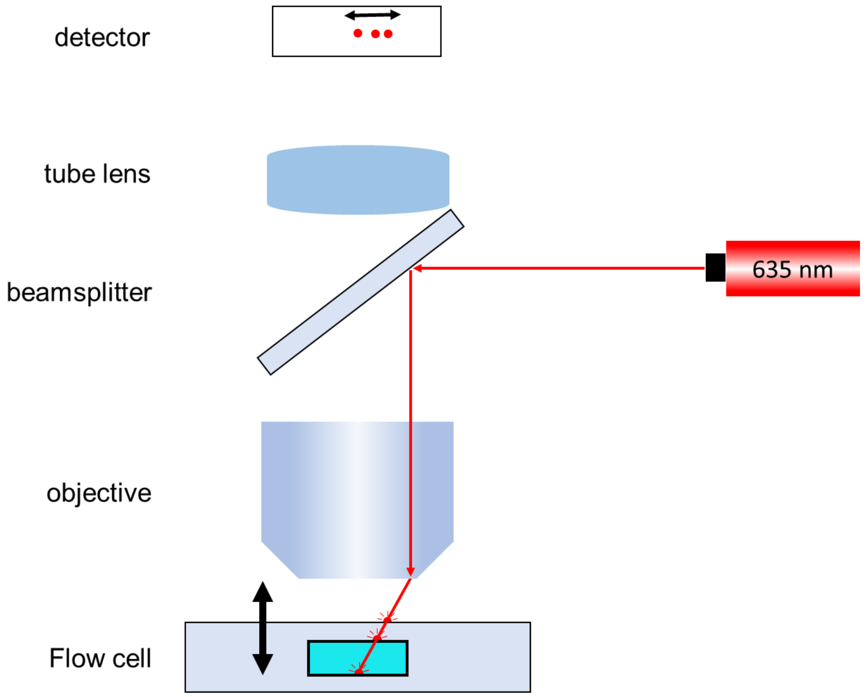


**Figure S1. Focus-finder optical configuration**. A 635 nm laser is collimated and directed into the system at an oblique angle to the imaging surface. The laser reflects off three distinct interfaces, producing multiple spots. As the objective is translated along the z-axis, these imaged laser spots shift laterally on the detector. The lateral position of each spot encodes the axial distance between the objective and the sample surface, enabling precise focus determination.

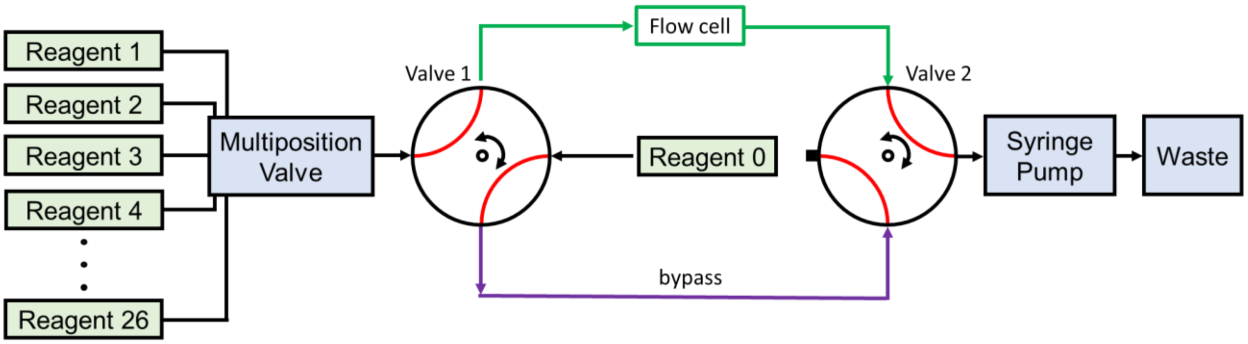


**Figure S2. Fluidics configuration.** Valves 1 and 2 can be configured in two positions, and together allow control over the source of the fluid (the multiposition valve or reagent 0) and the path (flow cell or flow cell bypass).

#### *i.c* MAPA Software

The hardware control was built from a baseline developed in MATLAB by the Greenleaf lab[1] that was first ported to MATLAB 2022 and subsequently modified. At the core of the system control lies a Windows 11 computer running MATLAB 2022. It controls all MAPA hardware components using appropriate control protocols for each: Kinetix Teledyne Camera via the Java version of Micromanager, and an external Arduino to handle illumination and camera trigger timing via serial, TC-36-25-RS232 Peltier temperature controller via a custom serial protocol, Kloehn PN24741 8 Channel V6 syringe pump via custom serial protocol, VICI EMHMA-CE selector valve via custom serial protocol, Labsmith uProcess automated 4-port selector valve AV202 via Python interface, KLD101 K-Cube laser diode driver via .NET interface, red and green lasers via combination of digital pins and custom serial protocol, and ASI MS-2000 XY stage and FW-1000 filter wheel via custom serial protocol.

The hardware control software runs an event loop that regularly checks in on all hardware components that have information to display at regular intervals (*e.g.*, lasers and their current measured output power levels, the Peltier element and its currently measured output power and temperature). This event loop runs constantly in the background. The collected data is displayed to a custom graphical user interface (GUI) for the instrument,which also allows for direct control over the stage position, temperature (ramping or instantaneous changes), laser power levels, camera exposure time, laser and filter settings, whether to snap an image, and fluidic controls (*e.g.* fluidic pathway, pumping speed and volume).

In addition, there is a separate GUI that allows the user to run protocols. First, the user can prime the lines to the sample tubes mounted on the instrument through a fluidic shortcut, bypassing the mounted flow cell. Next, they can define a protocol consisting of temperature ramps, fluidic pumping operations, and imaging steps. The protocol is defined in a Microsoft Excel file and subsequently gets read by the instrument. This workflow allows non-expert users to operate the instrument.

In detail, the different kinds control steps adhere to the following behavior: In steps where the temperature gets increased, the ramping speed is 6 ºC/min. For steps in which the temperature gets decreased, the ramping speed is 2 ºC/min to allow for slow annealing. In fluidic operations, the user can specify the flow speed and fluidic pathway (through or past the flow cell). In imaging steps, the instrument iterates over the tiles at which the Illumina MiSeq sequenced clusters on the flow cell. For every tile, it moves the flow cell section of interest under the objective of the microscope, then launches the autofocusing procedure, and—upon automatically finding the perfect focus position for that tile—captures an image with the programmed laser color, intensity, and exposure time.

**Autofocusing.** The autofocusing algorithm works by leveraging the scheme depicted in **Figure S1**. Each interface the laser passes through (*i.e.*, change in refractive index) causes a reflection. The first reflection occurs when the laser transitions from free space into the flow cell. This first spot is always visible. When the flow cell channel is immediately underneath the camera objective, there are two additional interfaces: from the glass into the liquid of the flow cell channel, and from the liquid of the flow cell channel into the glass again. This final reflection is the one we detect and use for autofocusing, as it corresponds to the interface we are interested in imaging. To detect the center of this third spot, we first apply simple thresholding to separate background pixels from the pixels comprising the spots created by the laser reflections. We also ignore pixels that are too close to the edges, as those are often spurious reflections from the sides of the flow cell channel. Next, we use connected component labeling to identify contiguous areas of bright pixels. From there, we merge connected components that are close enough (horizontally) to likely originate from the same spot. This leaves us with several sets of pixels, each of which we assume to belong to one spot. For each pixel in a set, we take its x/y coordinate and weight it by the brightness of the pixel. This gives us the weighted average of all the pixel locations, which we assume to be the center of the dot. As this is still a noisy measurement of a spot's center, we subsequently move the stage of the device vertically in intervals of 50 stage units and capture an estimate centroid location for each stage position. This results in a collection of stage z coordinates and their corresponding spot centroid x-coordinate locations. We then fit a straight line through the (z, x) pairs of points, resulting in an equation $z = f(x)$ that relates the z-coordinate of the stage with the x-coordinate of the centroid in the image. Through manual calibration, we found which x-coordinate $x^*$ of the centroid corresponds to a focused image. We feed this through $z = f(x)$ and arrive at an estimate for the optimal z-coordinate. We then repeat this process of sweeping the stage positions until two subsequent optimal z-coordinate estimates are sufficiently close to one another and declare the final estimate the focused z coordinate.

Because this algorithm relies on the flow cell being immediately underneath the objective to see the third spot/reflection, we find the center of the flow cell at the beginning of each experiment. To this end, we have the user move the objective over the flow cell once and subsequently run a binary search for the edges of the flow cell using the autofocusing laser. The binary search works as follows. We assume the initial x-coordinate to be over the flow cell channel, so any captured image will contain three dots. We then move approximately one flow cell channel width to the right, which guarantees that we are outside the flow cell channel. Images captured here will contain just the first reflection (*i.e.*, one dot). These are now our left and right bounds for where the edge of the flow cell may be. We then move to the middle of this range and capture an image. If it contains three dots, it must be within the flow cell and we move the left bound to this x-coordinate. If it contains less than three dots, we must still be outside the flow cell edge, and we move the right bound to this coordinate. We then repeat this process until the right edge of the flow cell is found and repeat it for the left side. For any future autofocusing attempt, we go to the middle of the flow cell, defined as the middle between the left and right sides.

**Event Loop.** MATLAB 2022 has a bug in which serial communication is interruptible during read and write operations. This cannot be prevented, because MATLAB does not have mutexes or semaphores due to being a strictly single-threaded language. Instead, we implemented a "mailbox" system, in which interrupting calls will read outputs intended for interrupted calls, place them in a "mailbox stack", and then finish its own communication. Any call whose communication got interrupted and intercepted will time out, and the system may then check the mailbox to retrieve the information that was intended for it.

**Illumination Control.** The laser shutter is controlled by a pin. When the pin switches state, it takes ~10 ms for the laser shutter to fully turn open/close. To ensure that our images are fully illuminated, we use an external Arduino Leonardo to turn on the laser 10 ms before triggering the camera's exposure. The camera has an output pin, which is high while the camera is exposing its sensor. The Arduino monitors this pin's state and keeps the laser shutter open for as long as the camera is still exposing. This way, the user can program the camera to have a different exposure time, and the Arduino will adaptthe exposure time automatically.

#### ***i.d*** Data Analysis

The outline of the image processing pipeline is shown in **Figure 1c**. From the perspective of our algorithm, we begin with the images from our own microscope system, detect peaks in the images, then align those peaks with the coordinates from the FastQ files generated by the Illumina MiSeq to locate where a given cluster with a given sequence is in each and every image, and finally extract the intensity at that location.

**Peak Detection.** For our peak detection algorithm, we first apply a band-pass filter to the image, to smooth out noise and remove the background. All pixels below a user-definable threshold are set to be zero. We then utilize an off-the-shelf scikit-learn implementation to find local maxima, which are defined as pixels that are brighter than any of their eight surrounding neighbors. For every image, this set of coordinates is saved in a file. This step takes ~3 s per image.

**Coordinate Alignment.** To align the FastQ coordinates from the Illumina MiSeq with the detected peaks in an image, we begin by loading the coordinates for the respective devices into RAM and correcting away the 2D distortion caused by the imaging system. We then multiply the MiSeq coordinate by a constant factor to transform it to the same length-scale as the image coordinates of our system.

We next begin with a rough alignment step. Here, we assume that the two sets of coordinates may be transformed into one another using pure x/y-translation. This assumption has held true in our experience thanks to our clamp design. To find a x/y translation that roughly aligns the two point clouds well, we first transform them into two images, where pixels containing a point are set to be 1, and pixels without a point are set to 0. We then blur these images to account for uncertainty in the true extracted position. Subsequently, we calculate the 2D correlation function of the two images. In case there is perfect alignment, there should be a peak at the location corresponding to this x/y translation. In practice, there may be spurious peaks in the 2D correlation function, so before detecting these peaks, we subtract away a version calculated on even more blurred artificial images. This may be thought of as calculating the correlation of the density functions of the two point clouds we are attempting to align, so the resultant 2D correlation function is that of spurious alignments. Subtracting this background away, we are left with a 2D correlation that is within a few pixels of the true alignment.

This first step only applies an x/y-transformation and assumes no rotation and scaling errors. We therefore proceed to a fine-tuning step. Here, we start with the transformation from the rough alignment as an initial guess and subsequently apply an algorithm, which is a modified version of the iterative closest-point algorithm. This iterative algorithm first applies the current best transformation to the MiSeq coordinates and calculates the eight closest neighboring points (within 30 px) from our imaging system. The hypothesis is that for most MiSeq coordinates, one of the eight closest neighbors is likely to be the correct assignment. All of the other (up to seven) neighboring points are spurious and random. For each pair of MiSeq and imaging coordinates, we calculate which x/y-translation would perfectly match these two points and aggregate these x/y-translations in a 2D histogram. As in the 2D correlation case, we expect a single peak stemming from the many correct pairs, and a diffuse background stemming from the spurious alignments. We find the true peak by applying a band-pass filter and finding the remaining peak. We then repeat this process, but instead of finding the x/y translation that makes two neighbors overlap, we find the combination of rotation and scale around a center that makes the pairs of points overlap. This sequence of x/y translation fine-tuning and rotation/scale transformation fine-tuning is repeated until no further improvements in the alignment scores can be found. As we can assume that the (up to seven) other neighboring points are random, we can model this random distribution and calculate the probability density function as a function of the peak density of our image. When we plot a histogram of the distance between each pair of MiSeq and imaging coordinates, we see this background distribution and—for good alignments—an additional peak. We subtract the spurious alignments from the histogram, leaving only the additional peak associated with the good alignments. We take the average of this peak to be the average mis-alignment of our current transformation in pixels. A good value is $< 1$ px. This step takes ~15 s/image.

**Camera Distortion Correction.** We model our 2D optical distortion using a simple Brown-Conrady even-order polynomial model. We define distortion as the transformation, which takes a "ground truth" coordinate and morphs it into the image coordinate system we observe. To undistort a coordinate (*i.e.*, transform from image coordinates to true coordinates), this model first subtracts a center coordinate from a given image coordinate and then applies a scaling factor, which is an even-order polynomial function of the radial distance from the center coordinate, and finally re-adds the center coordinate. To re-distort a coordinate, we use Newton's method to calculate the appropriate distortion to apply to each coordinate, which converges quickly as our distortions are quite small. To determine the coefficients of the even-order polynomial of our own imaging system, we took several overlapping images of the flow cell and tried aligning them. Through a parameter sweep, we found a set of distortion parameters that allowed for the best alignment of the two images. To determine the hidden coefficients of the even-order polynomial of the MiSeq system, we assumed the previously determined parameters of our own imaging system to be accurate, and used a parameter sweep to find parameters which result in the smallest error in alignment.

**Tile Coordinate Estimation.** In order to image a sequenced MiSeq tile, our imaging setup needs to know which stage x/y coordinate it should navigate to. This is important because the area covered by one image is only slightly larger than the size of one MiSeq tile. For this reason, we need to ensure that the images we capture are perfectly aligned with the MiSeq tile locations. We have found that it is a good assumption that our initial guess for the tile location does not change greatly between experiments thanks to our clamp design. It varies by at most ~1,600 px, or around half an image. Thus, there is always an overlap between the images we try to capture of a tile and the tile itself, allowing us to apply our coordinate alignment algorithm. We also store the x/y stage coordinates of where an image was captured. This, together with the information of how many stage coordinates correspond to a pixel—which may be calculated from the manual of the stage and calculating the size of a pixel from the specifications of the camera and the objective—is then used to estimate where the perfect x/y stage coordinate for a tile lies. We calibrate this at the start of each experiment.

**Movie Coordinate Alignment.** For movies, the flow cell never moves between frames. In this case, subsequent frames of the movie will usually be very close in terms of coordinate alignment. We significantly speed up the image alignment process by foregoing the initial rough alignment step and simply taking the final alignment from another, previously aligned frame as our initial guess.

**Intensity Extraction.** After finding the transformation that transforms the MiSeq coordinates to the image coordinates (*i.e.,* undistort MiSeq coordinates, apply orthogonal transformation, and reapply our imaging system's distortion), we then calculate the intensity of the area immediately around each MiSeq cluster. This extraction algorithm is valid, as our average error in alignment is typically <1 px. We first subtract out an estimate of the image's background signal by running a minimum filter with large kernel size and subsequently applying a low-pass filter i.e., we look for the darkest pixel within a 10 px region, which we assume to be pure background because we design our MiSeq flow cell density to be low enough to have gaps between our clusters, and then apply a low-pass filter to smooth out our estimated background. This background-subtracted image is then convolved with a 3 x 3 mask of ones, and we subsequently linearly interpolate the intensity value at each aligned MiSeq coordinate. This is equivalent to taking the sum of the 3 x 3 pixel area immediately surrounding the aligned MiSeq coordinate. This step takes ~4 s/image.

**Data Aggregation.** Having applied this data pipeline to every captured image, we then aggregate it into a Pandas data frame, containing an ID for each cluster, its sequence and Q-score, and its intensity value for every image that was captured.

#### *i.e* Library Preparation and High-Throughput Sequencing

To prepare the TWIST library for sequencing, we used the Illumina Nextera XT Index Kit v2 DNA. For the index PCR reaction, 0.25 pmol of library were mixed with 50 µL of each index N716 and S508, 250 µL of colorless GoTaq G2 Hot Start Master Mix, and nuclease-free water to a final volume of 500 µL. PCR was performed on an Eppendorf Mastercycler X50 using the following thermal cycling conditions: initial denaturation at 95 °C for 2 min, followed by 9 cycles of 95 °C for 30 s, 58 °C for 30 s, and 73 °C for 30 s. The PCR products were purified using the Axygen AxyPrep MAG PCR Clean-Up Kit (MAG-PCR-CL-50). The final double-stranded DNA was quantified using both a Qubit Fluorometer (Thermo Fisher Scientific) and a NanoDrop 2000 Spectrophotometer (Thermo Fisher Scientific) prior to sequencing. Libraries were sequenced on an Illumina MiSeq platform using the MiSeq Reagent Kit v3 (Illumina, Cat#. MS-102-3001).

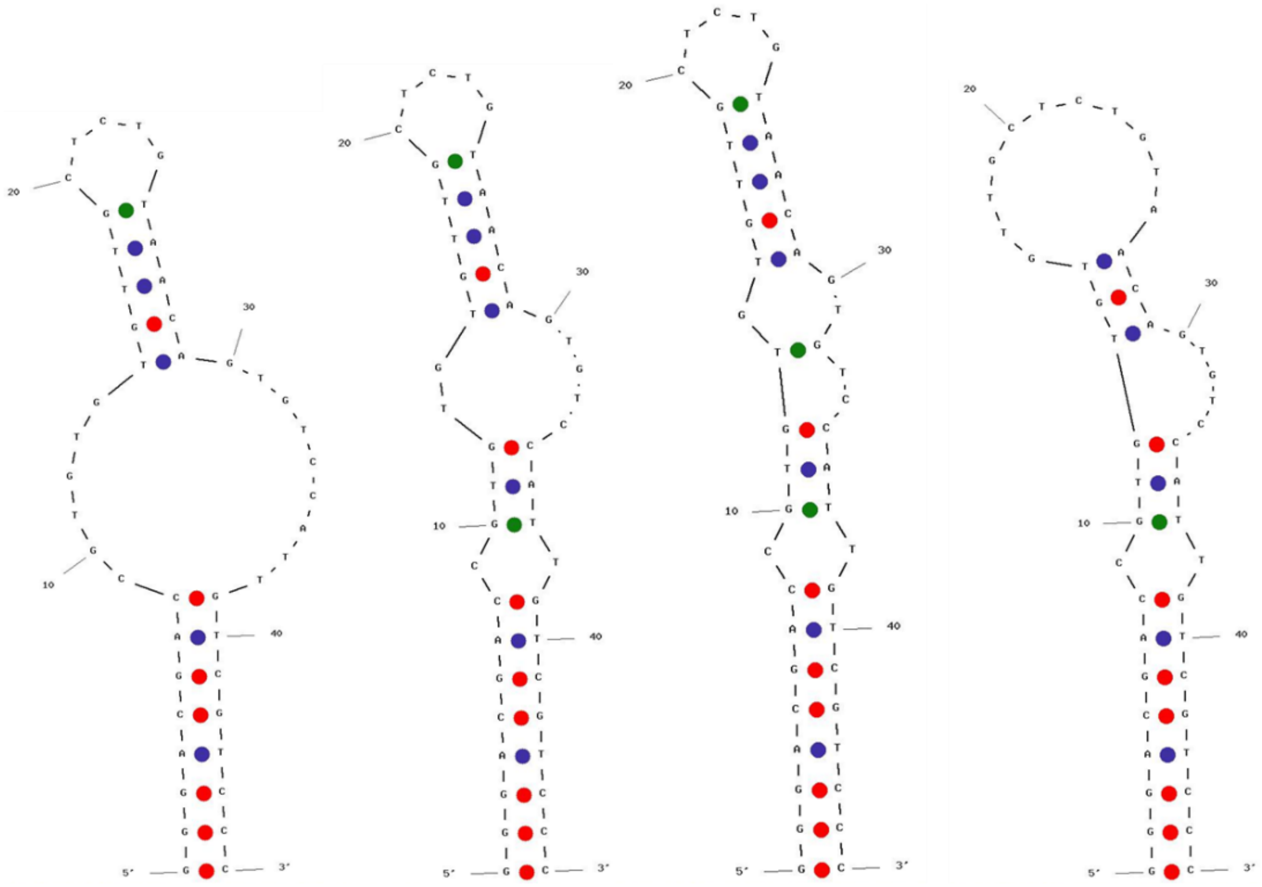


**Figure S3. MFold analysis of strand-displacement glucose aptamer.** The four most thermodynamically stable structures are shown, predicted using MFold at 25 °C, 1 M $Na^{+}$, and 10 mM $Mg^{2+}$.

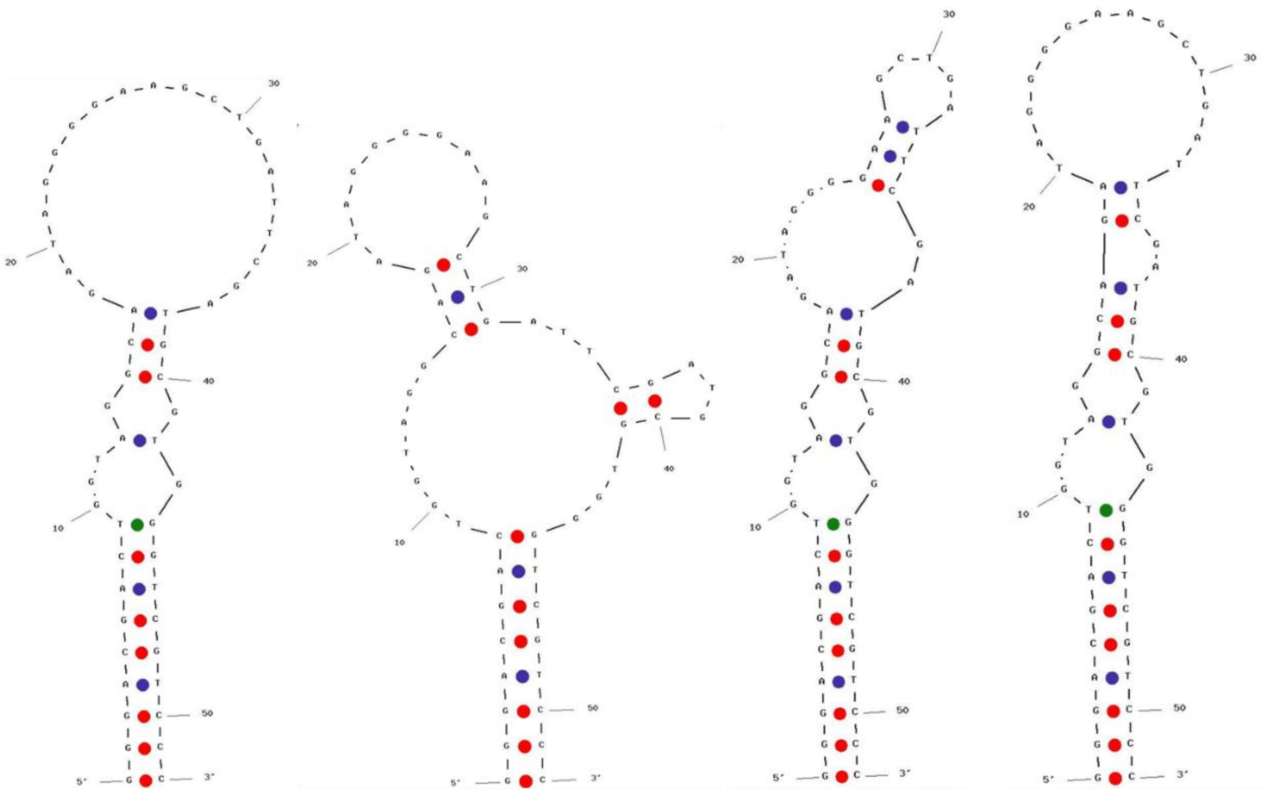

**Figure S4. MFold analysis of strand-displacement serotonin aptamer.** The four most thermodynamically stable structures are shown, predicted using MFold at 25 °C and 137 mM $Na^{+}$.

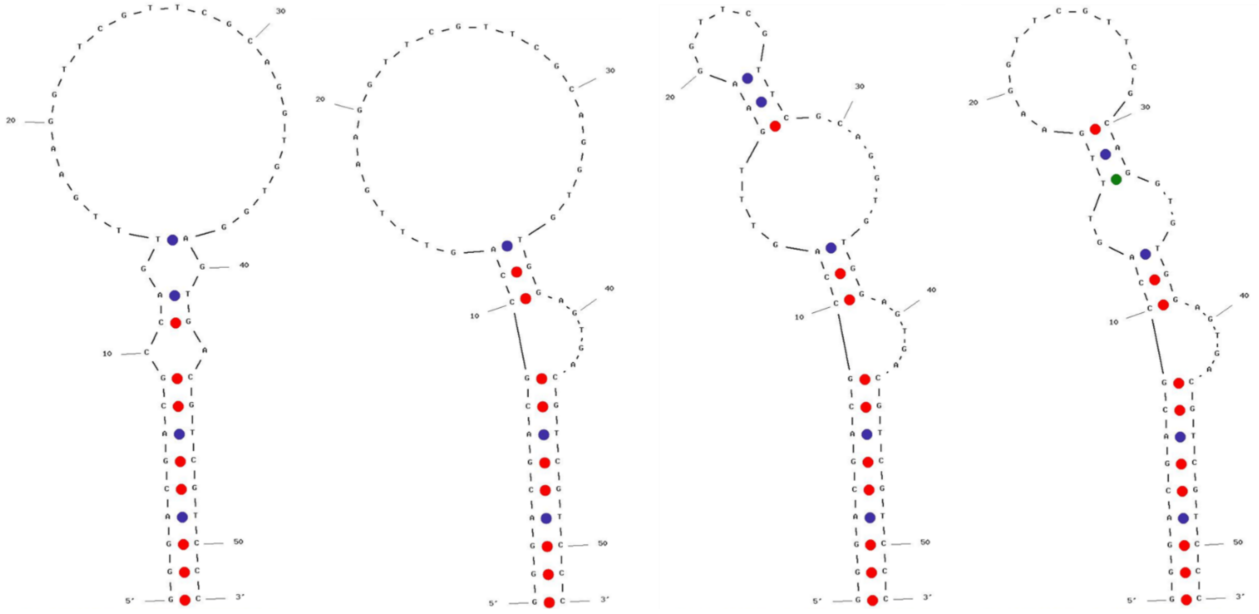

**Figure S5. MFold of strand-displacement dopamine aptamer.** The four most thermodynamically stable structures are shown, predicted using MFold at 25 °C and 137 mM $Na^+$.

### i.f Oligo Synthesis and Purification

**Solid-phase Synthesis of Oligonucleotides.** DNA was synthesized using an Applied Biosystems Expedite 8909 nucleic acid synthesis system. Native oligonucleotides employing standard β-cyanoethyl phosphoramidites as well as modified phosphoramidites were synthesized according to the manufacturer's instructions. In the case of Dabcyl CED phosphoramidite, we used 15 min incubation as specifications are not provided by the manufacturer. The final dimethoxytrityl (DMT) groups were retained after synthesis to facilitate purification using a Glen-Pak reversed-phase cartridge (Glen Research). Prior to purification, synthesized oligonucleotides were cleaved and deprotected with 30% ammonium hydroxide for 2 hours at 65 °C. Deprotected oligonucleotides were then purified using Glen-Pak reversed-phase cartridges following the provided protocol, and the resulting crude products were dried using a Thermo Fisher SpeedVac SRF110 refrigerated centrifugal vacuum concentrator.

**HPLC Purification of Oligonucleotides.** All synthesized oligonucleotides were purified using an Agilent 1260 Infinity II UHPLC equipped with either an Agilent PLRP-S reversed-phase HPLC column (300 Å, 3 µm, 4.6 mm × 150 mm, PL1512-3301) or a Phenomenex Luna reversed-phase C18 HPLC column (100 Å, 5 µm, 4.6 mm × 250 mm, 00G-4041-E0) The mobile phase consisted of 100 mM triethylammonium acetate (A) and acetonitrile (B).

For FAc-Dabcyl oligos, the gradient began with 18% B from t = 0 to 5 min, increased linearly to 23% B over 20 min, then quickly ramped to 95% B and was kept at 95% for 15 min. Mobile phase B was then quickly decreased to 18% and kept at 18% for 5 min. (Column: Luna). The flow rate was 1 mL/min, and the column was maintained at 50 °C.

For 5'FAM/3'Dabcyl oligos, the gradient began with 5% B from t = 0 to 5 min, increased linearly to 35% B over 20 min, then quickly ramped to 95% B, and was then held at 95% B from t = 25 to 40 min. Mobile phase B was then quickly decreased to 5% and kept at 5% for 5 min. (Column: PLRP-S RP). The flow rate was 1 mL/min, and the column was maintained at 60 °C.

Oligonucleotides were monitored by their intrinsic DNA absorbance at 260 nm, as well as by the characteristic absorbance of the attached labels Dabcyl (453 nm) and FAM (494 nm).

**Desalting of Oligonucleotides.** The HPLC-collected fractions were dried in a vacuum concentrator and resuspended in DI nuclease-free water. Finally, the eluent was desalted via size exclusion using a Glen Gel-Pak desalting column (Glen Research) and dried in a vacuum concentrator.

### i.g Oligo Sequences: Designed Oligos for MAPA Screening and Auxiliary Strands

**Table S1. Library Design.** Glucose (GLU), serotonin (SER), dopamine (DOPA), forward adaptor (FA), reverse complement of the reverse adaptor (RA),

| Name | Sequence |
|---|---|
| GLU_Twist | FA/GGGACGACCGTGTGTGTTGCTCTGTAACAGTGTCCATTGTCGTCC**CCATGG**/RA |
| SER_Twist | FA/GGGACGACTGGTAGGCAGATAGGGGAAGCTGATTCGATGCGTGGGTCGTCC**CCATGG**/RA |
| DOPA_Twist | FA/GGGACGACGCCAGTTTGAAGGTTCGTTCGCAGGTGTGGAGTGACGTCGTCC**CCATGG**/RA |
| FA | TCGTCGGCAGCGTCAGATGTGTATAAGAGACAG |
| RA | CTGTCTCTTATACACATCTCCGAGCCCACGAGAC |

**Table S2. Flow Cell Complementary Strands.** Adapter 1 complement (A1c), restriction site complement (RSc), extended lawn complements (ELc), Lawn complements (Lc), and quencher lawn complements (QLc) oligos.

| Name | Sequence |
|---|---|
| A1c-A550 | /5ATTO550N/CTGTCTCTTATACACATCTGACGCTGCCGACGA |
| A1c-Cy3 | /5Cy3N/CTGTCTCTTATACACATCTGACGCTGCCGACGA |
| A1c-Dabcyl | /Dabcyl/CTGTCTCTTATACACATCTGACGCTGCCGACGA |
| RSc-NCoI | /5Cy3/GTCTCGTGGGCTCGGAGATGTGTATAAGAGACAG**CCATGG**GGAC |
| EL1c | TTAGGGTTAGGGTTAGGGTTAGGGATCTCGTATGCCGTCTTCTGCTTG |
| EL2c | TTAGGGTTAGGGTTAGGGTTAGGGGTGTAGATCTCGGTGGTCGCCGTATCATT |
| L1c | ATCTCGTATGCCGTCTTCTGCTTG |
| L2c | GTGTAGATCTCGGTGGTCGCCGTATCATT |
| QL1c | /BHQ2/ATCTCGTATGCCGTCTTCTGCTTG |
| QL2c | /BHQ2/GTGTAGATCTCGGTGGTCGCCGTATCATT |

### i.h MAPA Molecular Protocols – Cluster Preprocessing and Functional Testing

Following sequencing, the MiSeq flow cell was mounted onto our MAPA platform for molecular preprocessing and high-throughput testing. To ensure proper optical coupling, 8 µL of refractive index-matched immersion liquid was applied between the flow cell and the prism. All steps were performed at 25 °C and all flow rates were 350 µL/min unless otherwise noted. Unless specified as a room temperature hybridization, all strand hybridizations were conducted as follows:

temperature was linearly ramped from 25–60 °C at a rate of 6 °C/min and held for 2 min. Subsequently, the temperature was reduced to 25 °C at 2 °C/min to allow hybridization.

**Flow Cell Pre-cleaning.** To remove unbound oligonucleotides, the flow cell was first incubated with formamide for 1 min, followed by a wash with PR2 buffer (provided by Illumina in the sequencing kit). To eliminate excess proteins, a detergent solution containing 1% SDS and 0.1% Triton X-100 in 1× PBS was then flowed through the flow cell, followed by another PR2 buffer wash.

**Hybridization for Alignment.** To match imaging positions with those used by the MiSeq, all clusters in the flow cell were fluorescently labeled by hybridizing at room temperature to a 100 nM Cy3-labeled strand complementary to the forward adapter (FAc-Cy3) in 1× SELEX buffer (20 mM Tris-HCl, 120 mM NaCl, 5 mM KCl, 1 mM $MgCl_2$, 1 mM $CaCl_2$, 0.01% Tween-20). Excess strands were removed by washing with 4 mL of PR2 buffer followed by 1 mL of SELEX buffer. A custom algorithm was then used to align images and generate a coordinates file for precise stage positioning during imaging.

**Lawns and Cluster Blocking.** To block the 3' end of oligonucleotides on the flow cell, we incubated the flow cell with a blocking solution containing: 1× TdT Reaction Buffer, 1 mM $CoCl_2$, 50 µM $ZnCl_2$, 1 mM ddTTP, 1 mM ddATP, and 4.8 U/µL TdT enzyme (Roche). The flow cell was incubated at 37 °C for 40 min, then washed with detergent solution followed by PR2 buffer. This process was repeated once.

**Cluster Cutting.** A complement strand solution containing 1 µM each of RSc-NcoI, L1c, and L2c in 1× Selex buffer was injected into the flow cell. The temperature was then ramped linearly to 60 °C at a rate of 6 °C/min and held for 2 min. Subsequently, the temperature was reduced to 25 °C at 2 °C/min to allow hybridization. Next, an NcoI solution (1 U/µL NcoI in 1× CutSmart buffer) was injected and incubated for 7 min at 37 °C. To remove excess protein, detergent solution was introduced and incubated for 3 min, followed by a wash with PR2 buffer. To remove hybridized DNA from the flow cell, formamide was flowed in and incubated for 1 min, followed by another PR2 wash. These three steps—complement strand hybridization, NcoI incubation, and washing—were repeated twice. Finally, the flow cell was washed with selection buffer.

**Cluster Labeling.** To minimize unintended labeling of residual unblocked lawn oligos (from the "Lawns and Cluster Blocking" step), a solution containing 1 μM each of EL1c and EL2c in 1× Selex buffer was injected into the flow cell. These complement strands are analogous to L1c and L2c but include a 24-mer overhang. This design takes advantage of the fact that TdT exhibits higher labeling efficiency on protruding 3′-OH ends compared to recessed or blunt ends.[2] To allow hybridization, the temperature was ramped linearly to 60 °C at a rate of 6 °C/min and held for 2 min. Subsequently, the temperature was reduced to 25 °C at 2 °C/min. To label the 3′ ends of cleaved clusters, the flow cell was then incubated at 37 °C for 40 min in a labeling solution containing 1× TdT Reaction Buffer, 1 mM $CoCl_2$, 50 μM 5-propargylamino-ddUTP-Cy3, and 1.5 U/μL TdT enzyme (NEB). Finally, the flow cell was washed sequentially with detergent solution followed by PR2 buffer.

**Quencher Strand Hybridization and Flow Cell Preparation.** A solution containing 100 nM FAc-Dabcyl, 1 μM QL1c, and 1 μM QL2c in the appropriate aptamer buffer (see below) was introduced into the flow cell for hybridization. The FAc-Dabcyl strand serves two purposes: it anneals to the adapter region to prevent possible interference with the aptamer domain by forming a duplex, and it positions the quencher near the base of the aptamer stem for signal modulation. QL1c and QL2c hybridize to the flow cell lawns to suppress background fluorescence from non-specific labeled lawns during cluster labeling with 5-propargylamino-ddUTP-Cy3.

**High-Throughput Screening of Aptamer Beacons.** After hybridization, the flow cell was washed with the appropriate aptamer buffer and incubated for 20 min. Clusters were then imaged to establish baseline fluorescence in the absence of target. To evaluate switch responsiveness and reversibility, the flow cell was subjected to cycles of target or buffer exposure followed by imaging. Each cycle consisted of: (1) target introduction and a 10-min incubation, (2) imaging, (3) buffer introduction and a 10-min incubation, and (4) a second imaging step. The final three cycles were performed using the highest target concentration. Target identities, concentrations, and buffer compositions are specified in the following section.

**Aptamer Buffers and Target Concentrations.** Glucose was tested at concentrations of 1 μM, 10 μM, 100 μM, 1 mM, 10 mM, and 100 mM. All experiments were performed in 1× HEPES buffer (20 mM HEPES, 1 M NaCl, 10 mM $MgCl_2$, 5 mM KCl, pH 7.4). Serotonin was tested at concentrations of 1 nM, 10 nM, 100 nM, 1 μM, 10 μM, 50 μM, and 100 μM. Experiments were conducted in 1× PBS buffer (10 mM $Na_2HPO_4$, 1.8 mM $KH_2PO_4$, 2.7 mM KCl, 137 mM NaCl). Dopamine was tested at concentrations of 1 nM, 10 nM, 100 nM, 1 μM, 10 μM, 50 μM, and 100 μM. Experiments were conducted in 1× PBS buffer.

#### i.i Measurement of Effective Binding Affinity via Fluorescence

To measure effective binding affinity, aptamers were labeled with 6-FAM at the 5′ end and Dabcyl at the 3′ end to enable fluorescence-based detection. Aptamer solutions were prepared at a concentration of 200 nM in the appropriate aptamer buffer. Target solutions were prepared separately in the same buffer, and 100 μL of each was mixed with 100 μL of the 200 nM aptamer solution. Samples were incubated for 10 min (for glucose and dopamine variants) and 12 h for serotonin variants. Fluorescence was measured in triplicate using a Qubit Fluorometer. All measurements were performed at room temperature, unless otherwise stated. Data were analyzed in Origin 2022 and fitted using the Hill equation with a fixed Hill coefficient (n = 1). In the model, $S$ and $T$ represent the observed fluorescence signal (in relative fluorescence units, RFU) and the target concentration, respectively. $B_{max}$ and $K_D$ are the fitted parameters:

$$S = S_o - B_{max} \frac{[T]^n}{K_D^n + [T]^n} \quad \text{(eq. S1)}$$

**i.j Raw Fluorescence Responses of the Top Four MAPA-identified Aptamer Beacons for Glucose, Serotonin, and Dopamine**

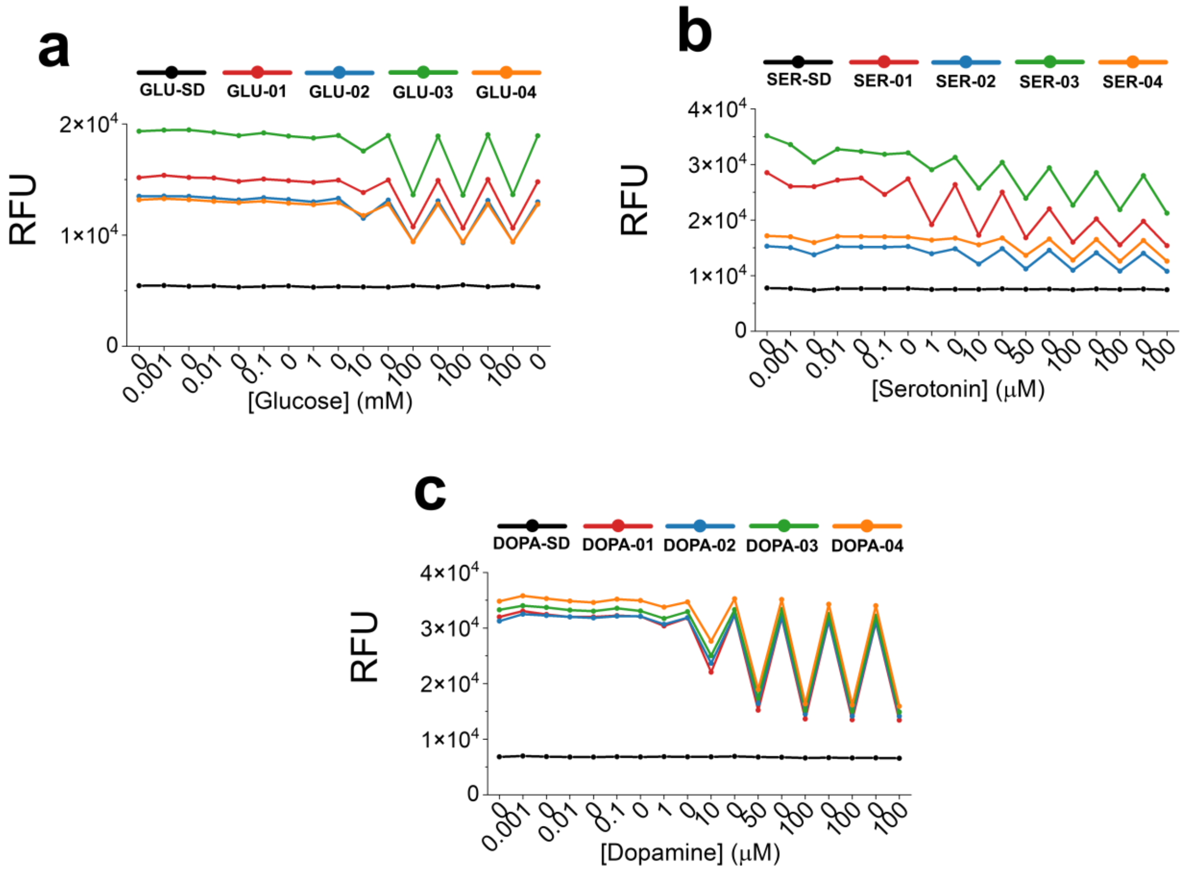


**Figure S7. Identification of high-performing aptamer beacons for glucose, serotonin, and dopamine using MAPA.** Raw fluorescence responses of the top four MAPA-identified aptamer beacons for glucose (**a**), serotonin (**b**), and dopamine (**c**), compared to their respective parental aptamers (GLU-SD, SER-SD, and DOPA-SD). The parental sequences show low signal and minimal or no response, consistent with their stable, quenched conformation. In contrast, the top-performing variants exhibit marked increases in signal upon target binding.
